## Supplemental figures for "Opto-RhoGEFs: an optimized optogenetic toolbox to reversibly control Rho GTPase activity on a global to subcellular scale, enabling precise control over vascular endothelial barrier strength"

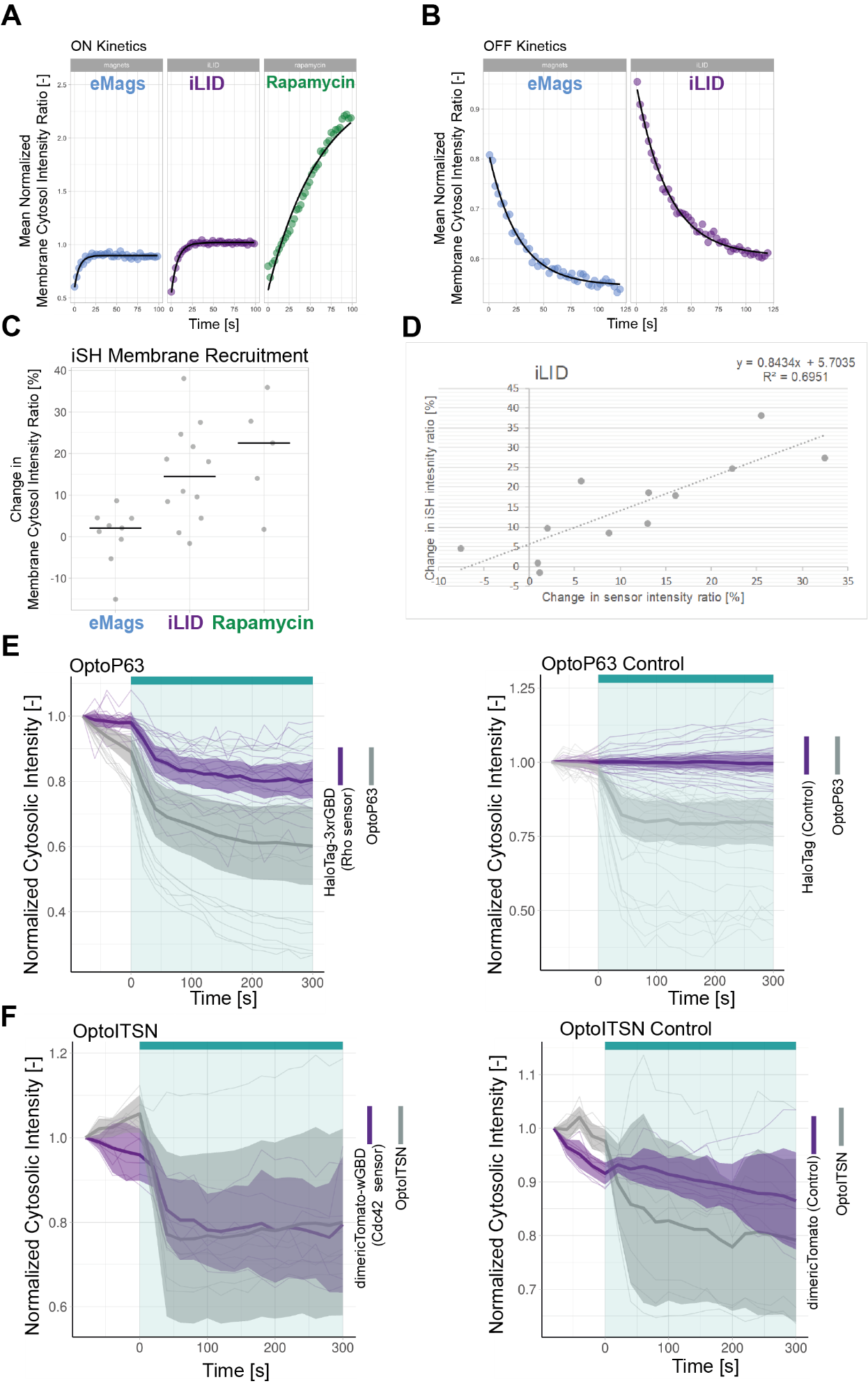

***Figure S1****.* ***Optogenetic Tool Setup***

*(****A****) Curve Fitting to determine t ½ ON kinetics shown in* ***Fig. 1A*** *for the mean normalized membrane to cytosol intensity ratio, where dots represent the mean values shown in* ***Fig. 1A****, blue = eMags, purple = iLID, green = rapamycin. The result of the curve fitting is sown as a black line. In the shown time range cells were activated with 488 nm laser light at 1 % laser power every 2.5 s or stimulated with 100 nM rapamycin at time point 0 s.(****B****) Curve Fitting to determine t ½ OFF kinetics for the normalized membrane to cytosol intensity ratio, where dots represent the mean values shown in* ***Fig. 1A****, blue = eMags, purple = iLID. The result of the curve fitting is sown as a black line. Time point 0 s is set to the first frame after photo activation. (****C****) Change in Membrane to Cytosol Intensity Ratio as percentage for the localization of iSH, localization of the PIP3 sensor is presented in* ***Fig. 1B****. All Hela cells were expressing the PIP3 location sensor mCherry-Akt-PH and for the iLID system Venus-iLID-CaaX, iSH-iRFP-SspB, for the eMags system* eMagA*-eGFP-CaaX and eMagB-iSH-iRFP670 and for the rapamycin system Lck-FRB-mTurquoise2 and mNeonGreen-FKBP12-iSH. Stimulated with either 100 nM rapamycin or 488 nm laser light each frame. Comparing the ratio of membrane over cytosol intensity of the iSH for 0 s pre activation and 50 s of activation. Each dot represents and individual cell. The median of the data is shown as a black bar. The number of cells per condition is: iLID = 29, eMags = 20, rapamycin = 15. (****D****) Plot of the correlation between change in iSH intensity ratio and change in PIP3 sensor intensity ratio for the data shown in* ***Fig. 1B, S1C****. The general linear model (top right corner) was fit to the data and is indicated as a dashed line. (****E****) Left: Normalized cytosolic intensity for the HaloTag-3xrGBD Rho sensor (purple) stained with JF635 nm dye and of the corresponding SspB-mCherry-p63RhoGEF(DH) (grey) intensity, upon photo activation (indicated by cyan bar) expressed in HeLa cells together with Lck-mTurquoise2-iLID. Showing the SspB-mCherry-p63RhoGEF(DH) recruitment for* ***Fig. 1F****. Thin lines represent individual cells; thick lines represent the mean values and ribbons represent their 95% confidence interval. The number of analyzed cells is 18. The data is from two biological replicates based on independent transfections. Right: The same settings and conditions apply except that the cells expressed the HaloTag as a control instead of HaloTag-3xrGBD Rho sensor. The number of analyzed cells is 30. (****F****) Left: Normalized cytosolic intensity for the dimericTomato-wGBD Cdc42 sensor (purple) and of the corresponding SspB-HaloTag-ITSN1(DHPH) (grey), stained with JF635 nm dye, intensity, upon the photo activation (indicated by cyan bar) expressed in HeLa cells together with Lck-mTurquoise2-iLID. Showing the SspB-HaloTag-ITSN1(DHPH) recruitment for* ***Fig. 1G****. Thin lines represent individual cells; thick lines represent the mean values and ribbons represent their 95% confidence interval. The number of analyzed cells is 6. The data is from two biological replicates based on independent transfections. Right: The same settings and conditions apply, except that the cells expressed the dimericTomato as a control instead of the dimericTomato-wGBD Cdc42 sensor. The number of analyzed cells is 30.*

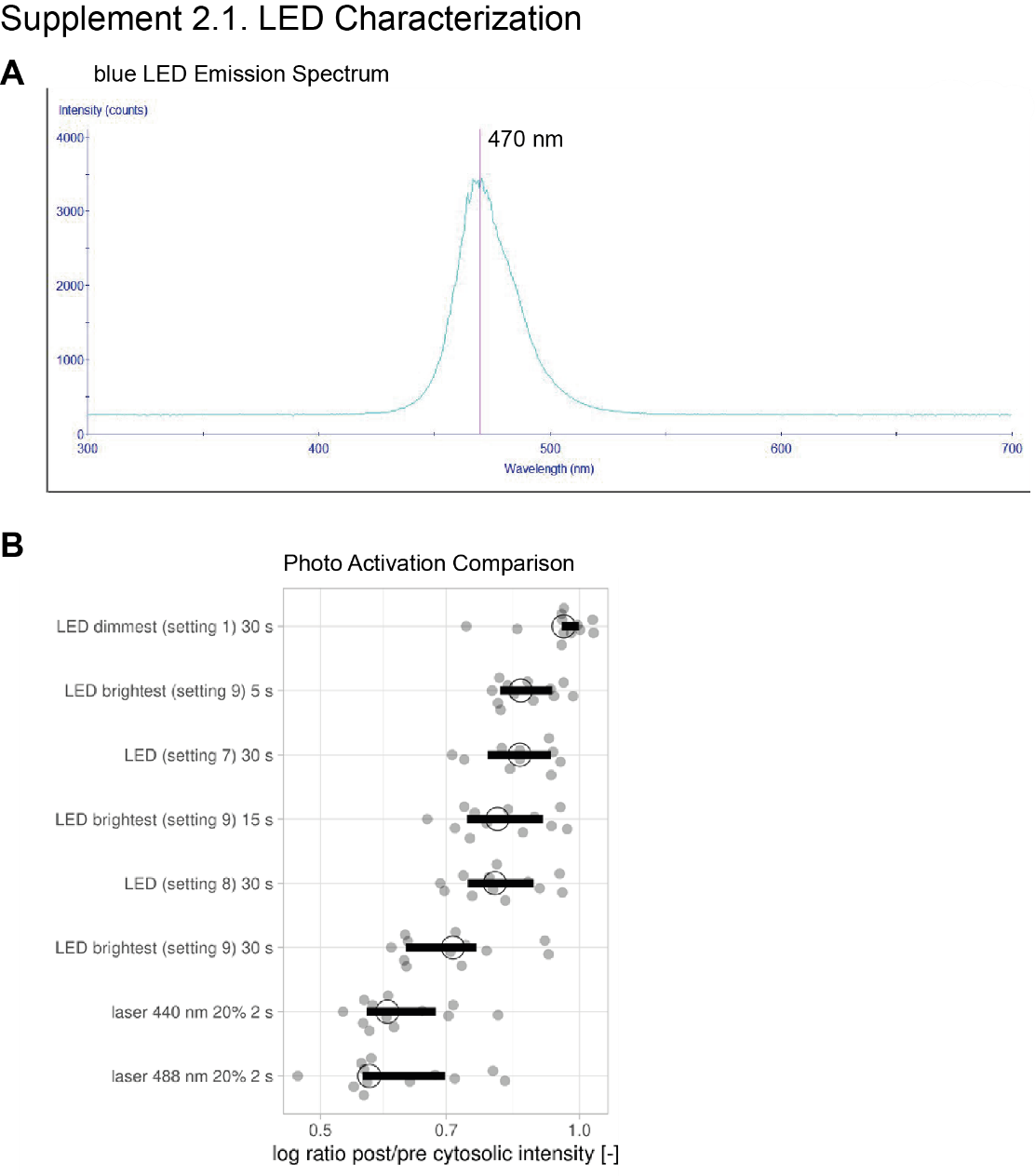

***Figure S2.1. Photo activation with blue LED.***

*(****A****) Emission spectrum of the LEDs in the blue setting. Red line indicates peak emission wavelength of 470 nm. (****B****) Recruitment efficiency as the log ratio of post over pre activation cytosolic intensity for HeLa cells expressing Lck-mTurquoise2-iLID and SspB-mScarlet-I activated with either blue LED light or laser light for the indicated time. Each dot represents and individual cell. The median of the data is shown as a black circle and the 95% confidence interval for each median, determined by bootstrapping, is indicated by the bar. The number of cells is 13, the same cells were imaged for the different photo activation settings.*

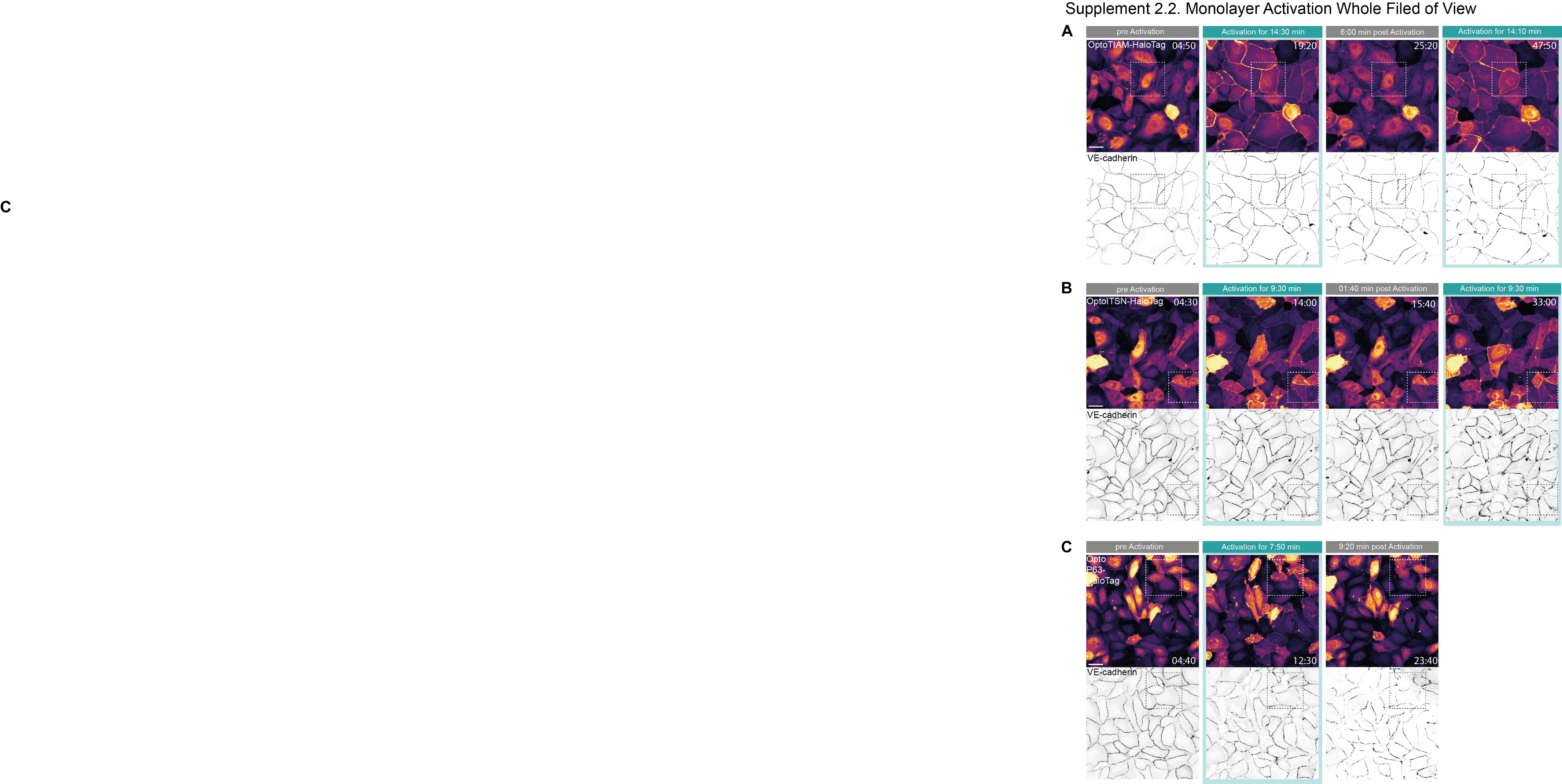

***Figure S2.2. Global photo-activation of endothelial cell monolayer***

*(****A****) Whole field of view confocal microscopy images of a BOEC monolayer stably expressing Lck-mTurquoise2-iLID (not shown) and either SspB-HaloTag-TIAM1(DHPH), ITSN1(DHPH) or p63RhoGEF(DH) stained with JF552 nm dye (LUT = mpl-inferno, brighter colors indicating higher intensity). Additionally, stained for VE-cadherin with the live labeling antibody Alexa Fluor 647 Mouse Anti-Human CD144 (gray inverted). Monolayer was photo activated with 442 nm laser light twice for 15 min with a 15 min dark recovery phase in between. Scale bars: 50 µm. Times are min:s from the start of the recording. Cyan bar indicates 442 nm photo activation. Dashed box indicated the zoom in shown in* ***Fig. 2C****. (****B****) The same settings and conditions apply as in* ***Fig. S2.2.A*** *but BOECs are expressing SspB-HaloTag-ITSN1(DHPH) and were photo activated twice for 10 min with 10 min dark recovery in between. (****C****) The same settings and conditions apply as in* ***Fig. S2.2.A*** *but BOECs are expressing SspB-HaloTag-p63RhoGEF(DH) and were photo activated once for 10 min with 10 min dark recovery afterwards.*

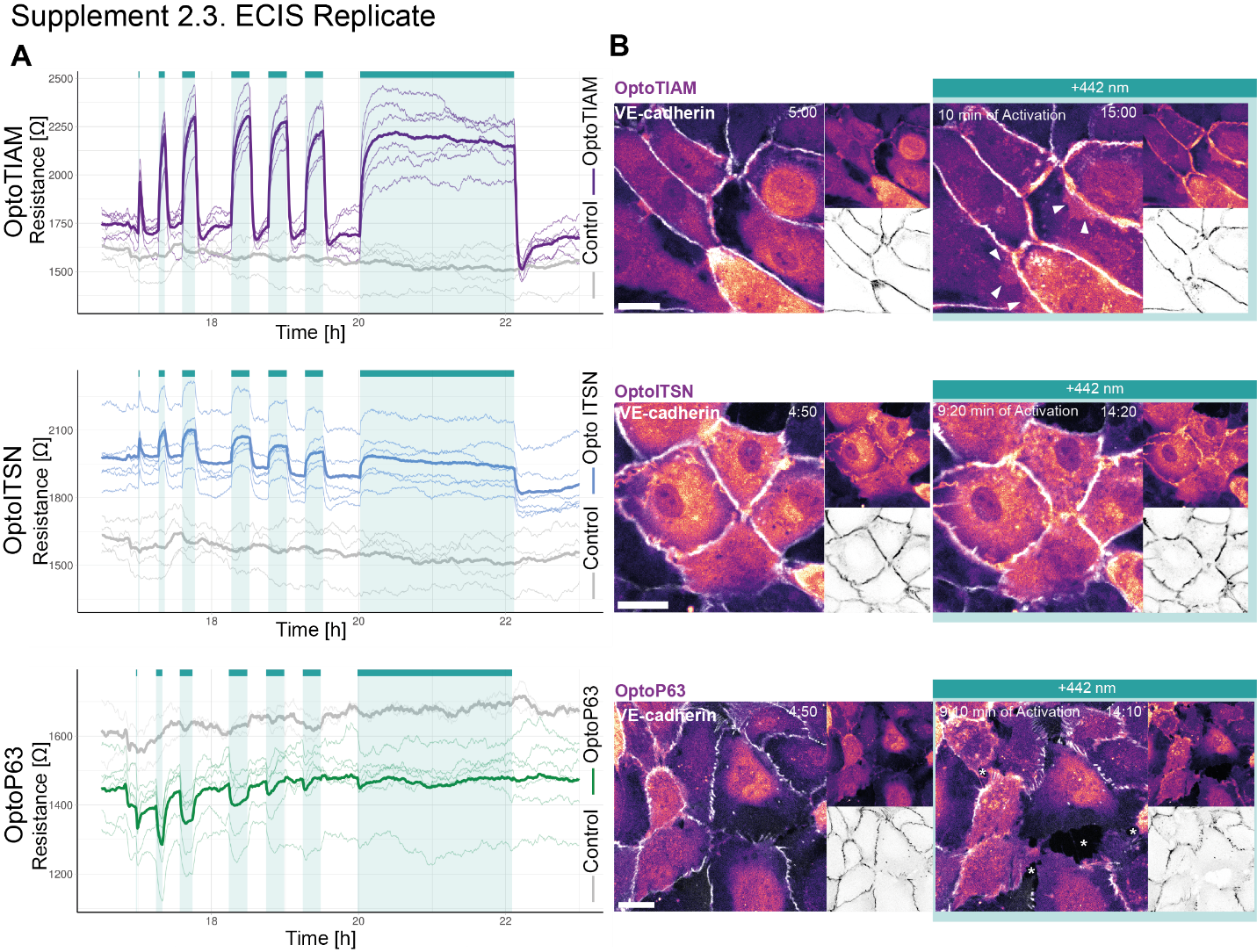

***Figure S2.3. Replicate ECIS assay and time-lapse microscopy***

*(****A****) Replicate for the resistance of a monolayer of BOECs stably expressing Lck-mTurquoise2-iLID, solely as a control (grey), and either SspB-HaloTag-TIAM1(DHPH)(purple)/ ITSN1(DHPH) (blue) or p63RhoGEF(DH) (green) measured with ECIS at 4000 Hz every 10 s. Cyan bars indicated photo activation with blue LED light (1 min, 5 min, 10 min, 3x 15 min, 120 min). Thin lines represent the average value from one well of an 8W10E ECIS array. Thick lines represent the mean. The number of wells per condition is: OptoTIAM=6, OptoITSN=6, OptoP63=6, control=4 and 2. (****B****) Representative zoom ins from confocal microscopy images of a BOEC monolayer stably expressing Lck-mTurquoise2-iLID (not shown) and either SspB-HaloTag-TIAM1(DHPH)/ ITSN1(DHPH) or p63RhoGEF(DH) stained with JF552 nm dye (LUT = mpl-magma, bright colors indicating higher intensity). Additionally, stained for VE-Cadherin with the live labeling antibody Alexa Fluor 647 Mouse Anti-Human CD144 (white in merge, grey inverted in single channel). Scale bars: 25 µm. Times are min:s from the start of the recording. Cyan bar indicates 442 nm photo activation. Arrows indicate overlap and protrusions. Asterisks indicate holes in monolayer.*

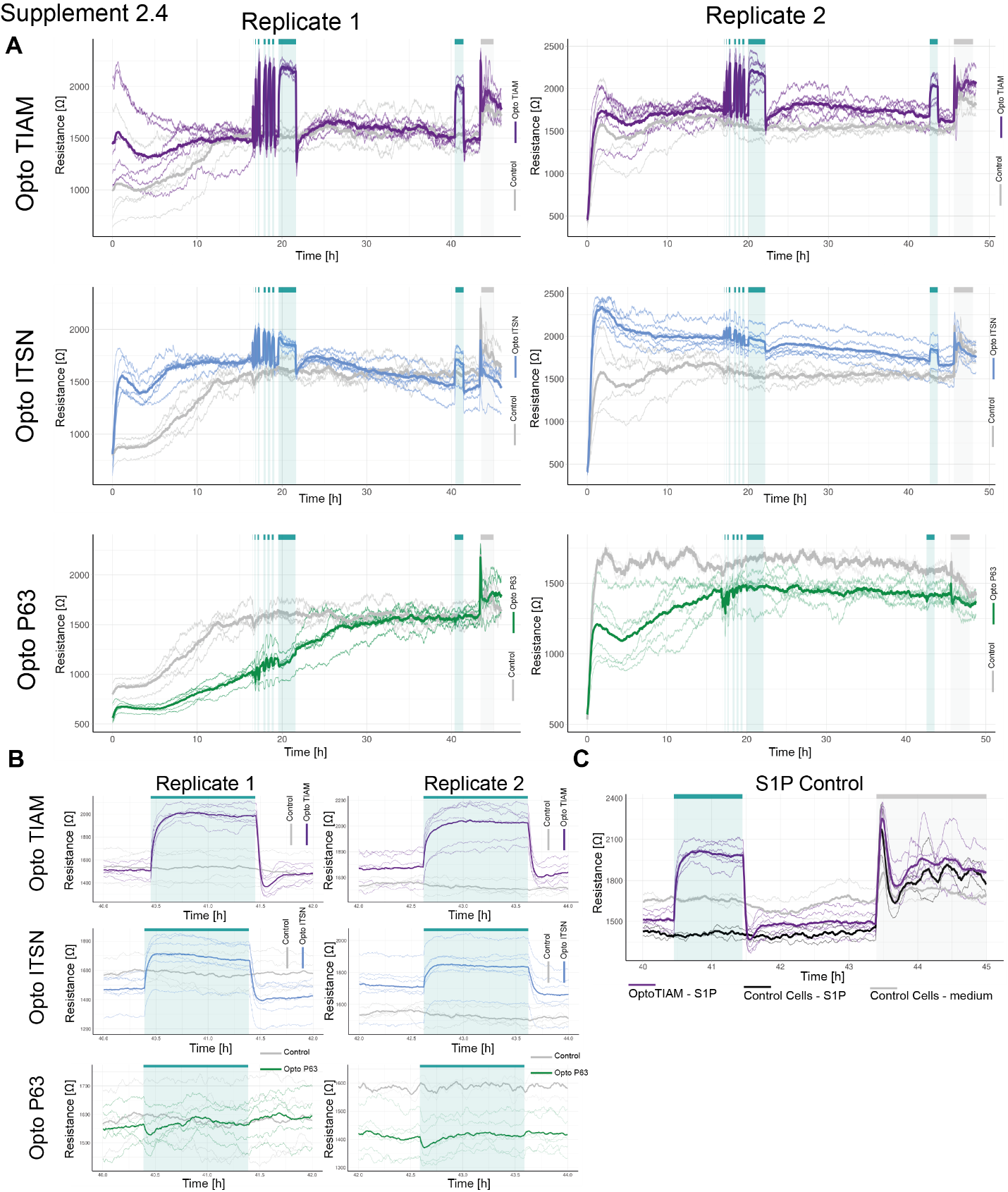

***Figure S2.4.*** **Overview entire time course ECIS assay**

*(****A****) ECIS experiment overview for the resistance of a monolayer of BOECs stably expressing Lck-mTurquoise2-iLID, solely as a control (grey), and either SspB-HaloTag-TIAM1(DHPH)(purple)/ ITSN1(DHPH) (blue) or p63RhoGEF(DH) (green) measured with ECIS at 4000 Hz every 10 s. Cyan bars indicated photo activation with blue LED light (1 min, 5 min, 10 min, 3x 15 min, 120 min, 60 min). Grey bar indicates S1P stimulation. Thin lines represent the average value from one well of an 8W10E ECIS array. Thick lines represent the mean. The number of wells per condition is: OptoTIAM=6, OptoITSN=6, OptoP63=6, control=4. The experiment was performed twice on different days. (****B****) Zoom in on the 1 h photo activation on the second day after seeding. The same settings and conditions apply as in* ***A****. (****C****) Zoom in on S1P stimulation for BOECs stably expressing Lck-mTurquoise2-iLID, solely as a control (grey), and either SspB-HaloTag-TIAM1(DHPH)(purple) for replicate 1. Cell were stimulated with 650 nM S1P in medium, OptoTIAM-S1P (purple) n=6 and Control-cells-S1P n=2 (black), as a control for the stimulation, cells were treated with medium only: Control-Cells –medium (grey) n=2.*

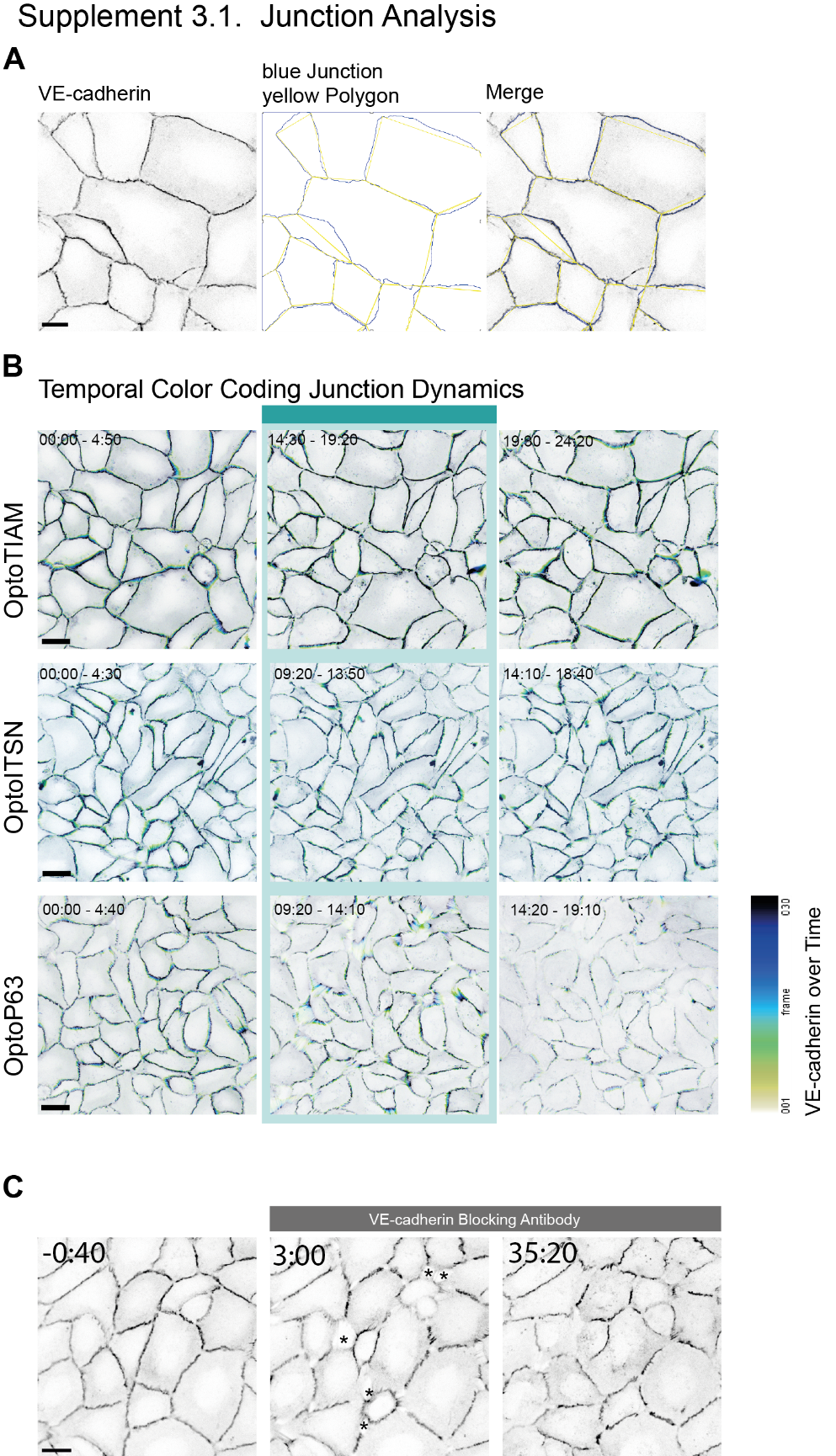

***Figure S3.1. Junction Analysis***

*(****A****) Left panel: Confocal microscopy image of a BOEC monolayer stably expressing Lck-mTurquoise2-iLID (not shown) and SspB-HaloTag-TIAM1(DHPH) (not shown), additionally, stained for VE-Cadherin with the live labeling antibody Alexa Fluor 647 Mouse Anti-Human CD144 (gray inverted). For the last frame before photo activation (pre). Middle panel: Blue lines represent the in Tissue Analyzer created junction outlines. Yellow lines represent the hand drawn polygon to measure the junction linearity index. Right panel: Shows a merge of the microscopy image and the lines used for the linearity index measurements. Scale bar: 25 µm (****B****) Temporal color coding for confocal microscopy images of VE-cadherin stained with the live labeling antibody Alexa Fluor 647 Mouse Anti-Human CD144, for a monolayer of BOECs stably expressing Lck-mTurquoise2-iLID (not shown) and either SspB-HaloTag-TIAM1(DHPH)/ ITSN1(DHPH) or p63RhoGEF(DH) (not shown). For the time period before photo activation (left panel), during photo activation indicated with cyan bar (middle panel) and after photo activation (right panel). Color code indicates time shown in look up table. Times are min:s from the start of the recording. Scale bars:50 µm. These images are also shown in* ***Fig.*** ***S2.2*** *and in* ***Fig. 2C*** *(****C****) Confocal microscopy image of a BOEC monolayer stably expressing Lck-mTurquoise2-iLID (not shown) and SspB-HaloTag-TIAM1(DHPH) (not shown), additionally, stained for VE-Cadherin with the live labeling antibody Alexa Fluor 647 Mouse Anti-Human CD144 (gray inverted). Cells were treated with VE-cadherin blocking antibody at time point zero, indicated by gray bar. Times are min:s starting from the treatment. Scale bar: 25 µm. Asterisks indicate holes in monolayer.*

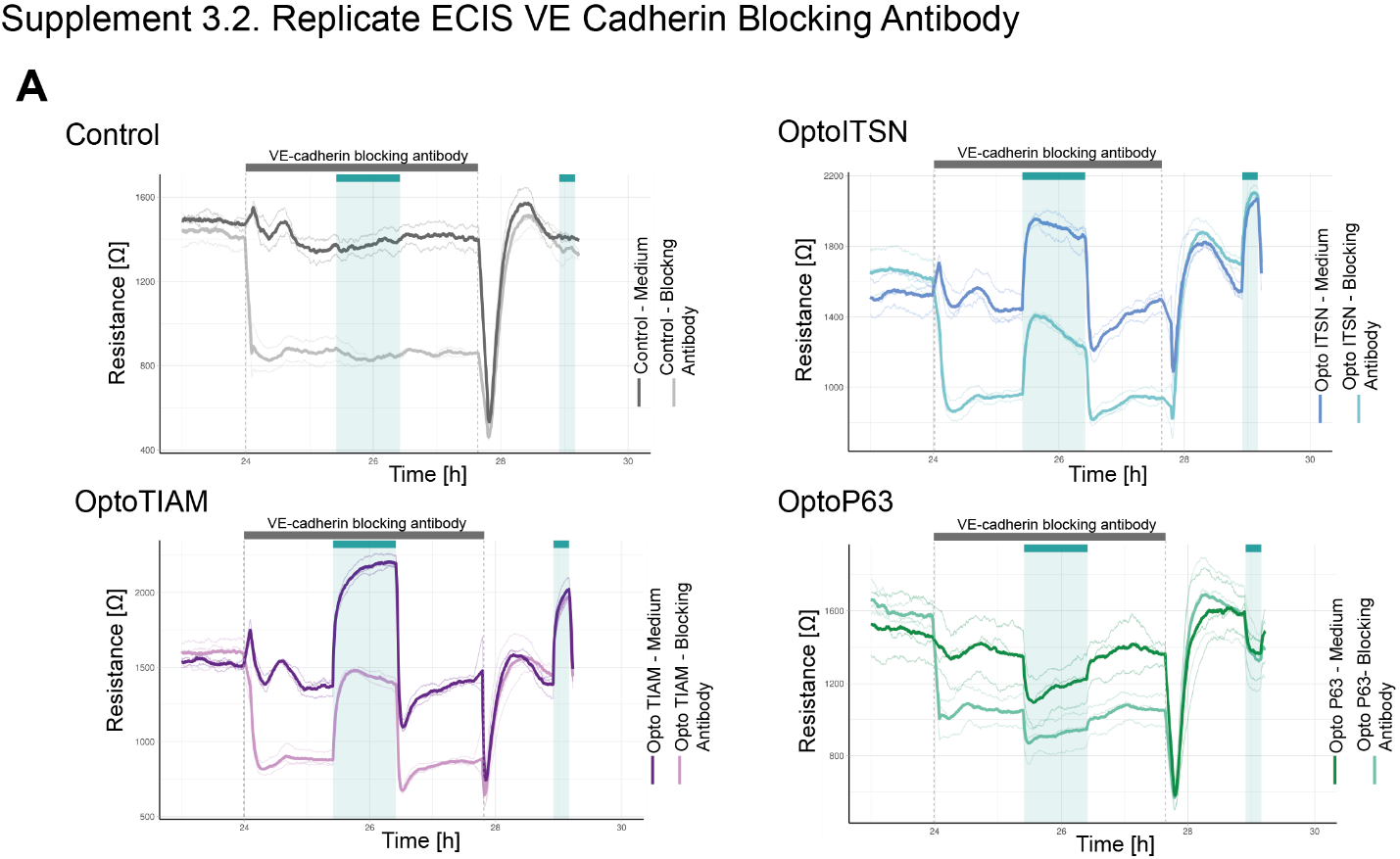

***Figure S3.2. Replicate ECIS assay in presents of the VE-cadherin blocking antibody***

*(****A****) Replicate for the resistance of a monolayer of BOECs stably expressing Lck-mTurquoise2-iLID, solely as a control (grey), and either SspB-HaloTag-TIAM1(DHPH)(purple)/ ITSN1(DHPH) (blue) or p63RhoGEF(DH) (green) measured with ECIS at 4000 Hz every 10 s. Cyan bars indicated photo activation with blue LED light (60 min, 15 min). Gray bar with dashed lines indicates the addition of VE-cadherin blocking antibody in medium (darker color line) or medium as a control (lighter color line). At the end of the gray bar the medium is replaced for all conditions. Thin lines represent the average value from one well of an 8W10E PET ECIS array. Thick lines represent the mean. Three wells were measured for each condition.*

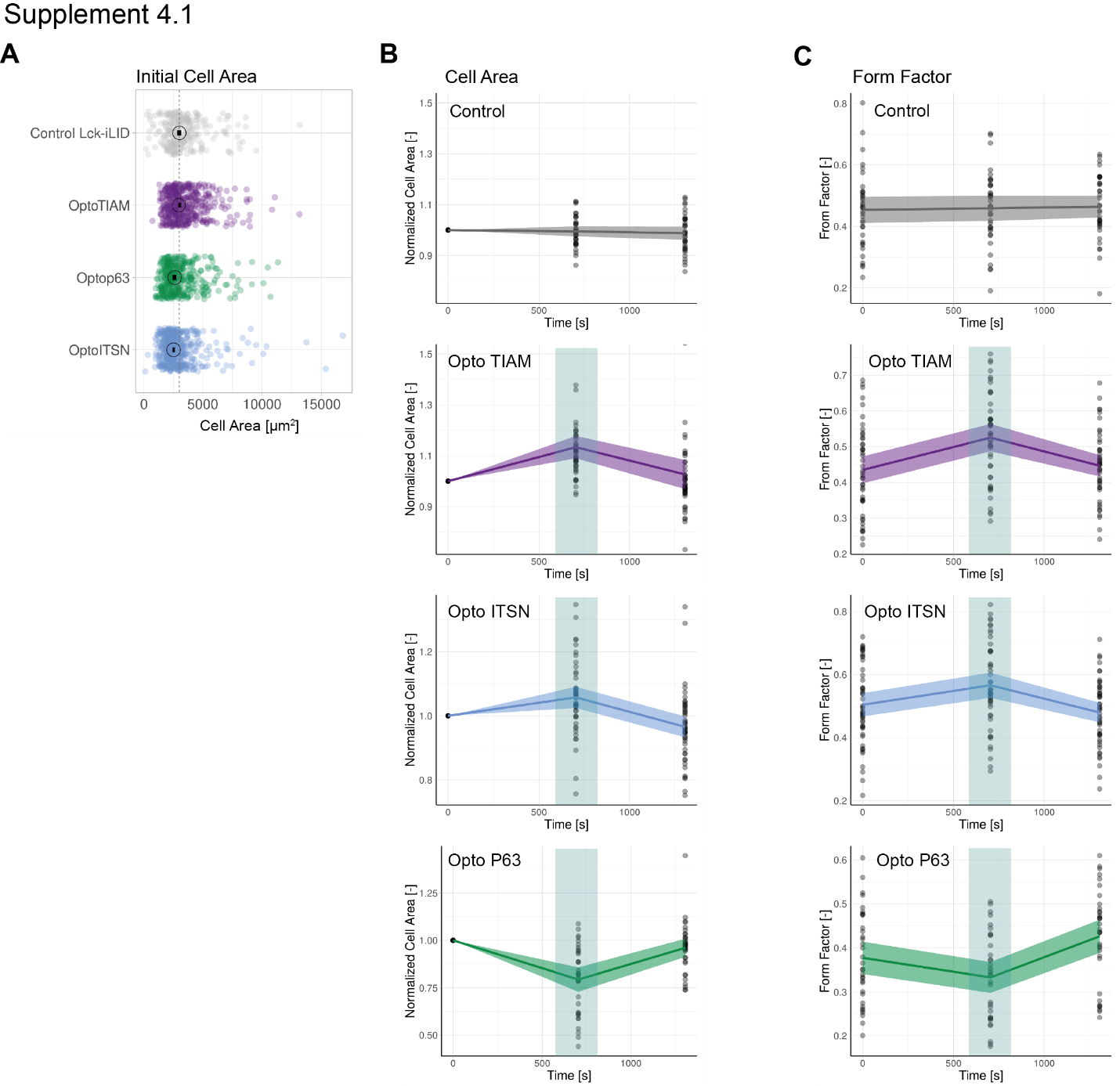

***Figure S4.1****.* ***Global photo activation***

*(****A****) Initial cell area for BOECs stably expressing Lck-mTurquoise2-iLID, solely as control, and either SspB-HaloTag-TIAM1(DHPH)/ ITSN1(DHPH) or p63RhoGEF(DH). Each dot represents and individual cell. The median of the data is shown as a black circle and the 95% confidence interval for each median, determined by bootstrapping, is indicated by the bar. The dashed line represents the median of the control condition. The number of cells per condition is: Control Lck-iLID=257, OptoITSN=467, Optop63=389, OptoTIAM=458. The data is from two independent experiments. (****B****) Cell area for BOECs stably expressing Lck-mTurquoise2-iLID, solely as control, and either SspB-HaloTag-TIAM1(DHPH)/ ITSN1(DHPH) or p63RhoGEF(DH) before activation (t=0s) at the end of 10 min activation (t=705) and 10 min after this activation (t=1305). Including the cell shown in* ***Fig. 4A,B****. Dots represent individual cells, thick lines represent the mean values and ribbons represent their 95% confidence interval. The number of analyzed cells is: Control=36, Opto-TIAM=45, Opto-ITSN=48, Opto-P63=34. The data is from two independent experiments with at least four individual photo-activations. (****C****) Form Factor for the in* ***B*** *described dataset and conditions.*

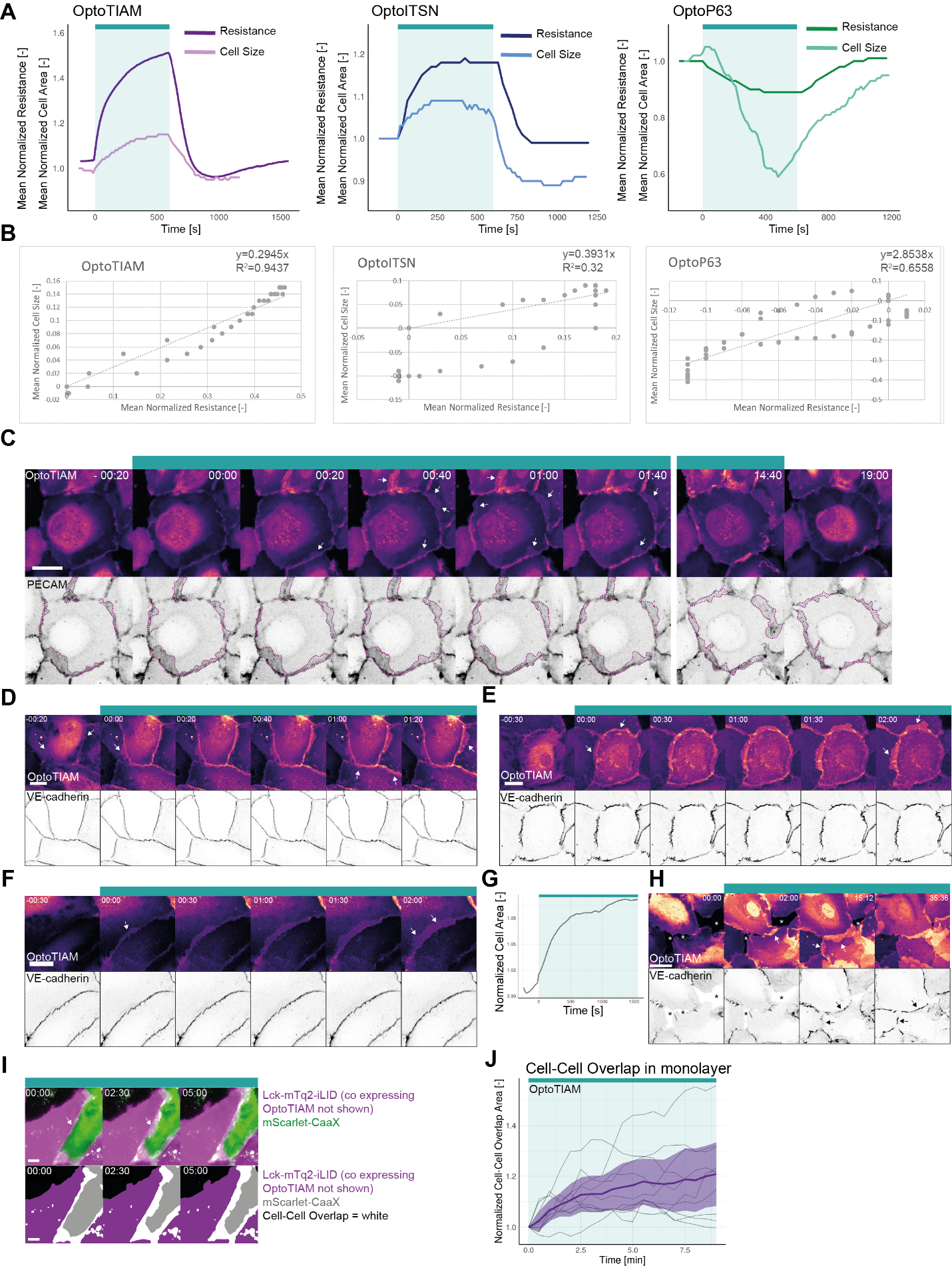

***Figure S4.2. Cell-cell overlap***

*(****A****) Normalized mean resistance and cell size for a 10 min photo-activation, indicated with cyan bar, of BOECs expressing Lck-mTurquoise2-iLID and either SspB-HaloTag-TIAM1(DHPH)(purple)/ ITSN1(DHPH) (blue) or p63RhoGEF(DH) (green). Resistance measured with ECIS (darker color) and cell size measured in confocal microscopy images (lighter color). The same data is also presented in* ***Fig. 2B*** *and* ***Fig 4B****. (****B****) Plot of the correlation between normalized mean resistance and normalized mean cell size for the curves shown in* ***Fig. S4.2A****. The general linear model (top right corner) was fit to the data and is indicated as a dashed line. (****C****) Confocal microscopy images from a time laps of a BOEC monolayer stably expressing Lck-mTurquoise2-iLID (not shown) and SspB-HaloTag-TIAM1(DHPH) (upper panel, LUT=mpl-magma, brighter colors represent higher intensity), additionally, stained for PECAM with a live labeling antibody Alexa Fluor® 647 Mouse Anti-Human CD31 (lower panel, inverted grey). Purple dashed line indicates higher intensity PECAM areas marking cell-cell overlap. Arrows indicate higher intensity in the Opto-TIAM channel in cell-cell overlap areas. Time is min: s from the start of photo-activation. The cyan bar indicates photo-activation with 442 nm laser light for 15 min. Scale bar: 25 µm. (****D, E, F****) Confocal microscopy images from a time laps of a BOEC monolayer stably expressing Lck-mTurquoise2-iLID (not shown) and SspB-HaloTag-TIAM1(DHPH) (upper panel, LUT=mpl-magma, brighter colors represent higher intensity), additionally, stained for VE-cadherin with the live labeling antibody Alexa Fluor 647 Mouse Anti-Human CD144 (lower panel, inverted grey). Arrows indicate higher intensity in the Opto-TIAM channel in cell-cell overlap areas. Time is min: s from the start of photo-activation. The cyan bar indicates photo-activation with 442 nm laser light. Scale bar: 25 µm. Cell E is also shown in* ***Fig. 2C****. (****G****) Normalized cell area over time for a sub-confluent monolayer of BOEC stably expressing Lck-mTurquoise2-iLID and SspB-HaloTag-TIAM1(DHPH), photo-activated with 442 nm laser light in the time period indicated with the cyan bar. (****H****) Zoom in on the corresponding confocal microscopy images measured in* ***Fig. S4.2G.*** *Images show BOECs, in a sub-confluent monolayer, stably expressing Lck-mTurquoise2-iLID (not shown) and SspB-HaloTag-TIAM1(DHPH) (upper panel, LUT=mpl-magma, brighter colors represent higher intensity), additionally, stained for VE-cadherin with the live labeling antibody Alexa Fluor 647 Mouse Anti-Human CD144 (lower panel, inverted grey). Asterisks indicate holes in the monolayer. White arrows indicate overlap. Black arrows indicate newly formed adherens junctions. Time is min: s from the start of the recording. The cyan bar indicates photo-activation with 442 nm laser light. Scale bar: 25 µm. (****I****) Wide field microscopy images of a mosaic BOEC monolayer with one population stably expressing SspB-HaloTag-TIAM1(DHPH) (not shown) and Lck-mTurquoise2-iLID, and the other population stably expressing the membrane marker mScarlet-CaaX. The cell-cell overlap is represented by the merge (top panel) of the Lck-mTurquoise2-iLID (magenta) and mScarlet-CaaX (green) channel and as a binary image (bottom panel) representing the cell area of Lck-mTurquoise2-iLID (purple) and mScarlet-CaaX (grey) cells, where the overlap is colored white. Cells were photo-activated with 442 nm excitation light as indicated by the cyan bar. Arrows indicate cell-cell overlap. (****J****) Quantification of the cell-cell overlap are for the in* ***I*** *presented experiment. Thin lines represent individual cells; the thick line represents the mean values and ribbons represent the 95% confidence interval. The number of analyzed cells is 10 for 3 individual photo-activations.*

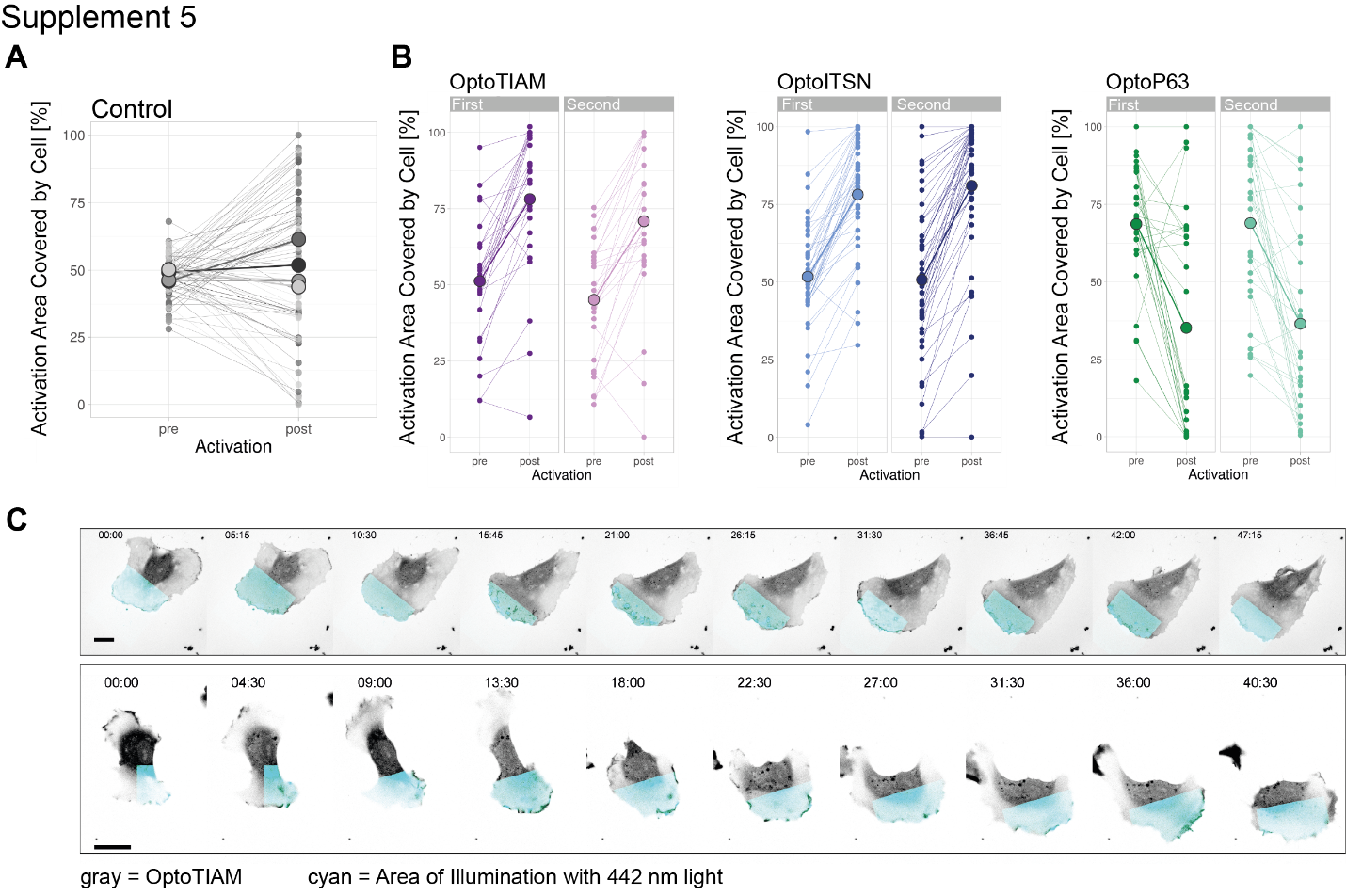

***Figure S5. Local Photo-activation***

*(****A****) Activation area covered by cell before and after 10 min without local photo-activation for control BOECs stably expressing Lck-mTurquoise2-iLID. Small dots represent individual areas, which are connected by lines. Larger dots represent the mean of each replicate indicated by different colors. The number of areas is: control=76. (****B****) Activation area covered by cell, pre and post 10 min activation with 442 nm light, for BOECs stably expressing Lck-mTurquoise2-iLID and either SspB-HaloTag-TIAM1(DHPH)(purple), -ITSN1(DHPH) (blue) or -p63RhoGEF(DH) (green), shown in* ***Fig. 5B****. Separated for first and second 10 min activation on the same cell but on opposite sides of the cell. Small dots represent individual activation areas, which are connected by lines. Larger dots represent the mean. The number of activation areas for Opto-TIAM=53, for Opto-ITSN=85, Opto-P63=61. (****C****) Stills from a confocal time lapse of a BOEC stably expressing Lck-mTurquoise2-iLID (not shown) and SspB-HaloTag-TIAM1(DHPH) (grey) stained with JF552 nm dye, locally photo-activated with 442 nm laser light in the blue area. Scale bars: 25 µm. Times are min:s from the start of the recording.*
